# Claustral Input Consolidates Anterior Cingulate Cortical States for Selective Behavior

**DOI:** 10.64898/2026.01.07.698269

**Authors:** Noa Peretz-Rivlin, Yonatan Fatal, Sivan Levin, Omri Gold, Keren Profesorsky, Maya Groysman, Ami Citri

**Affiliations:** The Hebrew University; The Hebrew University of Jerusalem

## Abstract

Efficient behavior depends on consolidating selective strategies that prioritize informative cues and suppress irrelevant signals. Previous studies have implicated the claustrum in coordinating cortical activity and large-scale brain states across behavioral contexts, yet its precise contribution to online behavioral control versus the consolidation of frontal control strategies remains unresolved. Here we combined large-scale Neuropixels recordings with targeted optogenetic perturbation of claustral neurons projecting to the anterior cingulate cortex in behaving mice. During adaptation to altered task context and constraints, frontal cortical activity reorganized into a coordinated control regime characterized by sharpened encoding of task-relevant cues, suppression of non-instructive signals, structured low-frequency dynamics, and rapid transitions between population-level neural states. Temporally misaligned claustral signaling disrupted this reorganization, selectively impairing the consolidation of efficient, cue-selective strategies across sessions without affecting trial-by-trial task execution. These findings identify a role for the claustrum in shaping frontal dynamics that consolidate adaptive control strategies.

## Introduction

Adaptive behavior depends on the ability to attain efficient strategies that balance reward and effort, maximizing success while minimizing unnecessary cognitive expenditure (Kolling et al., 2016; Shenhav et al., 2013; Walton & Bouret, 2019). This trade-off promotes cognitive economy, whereby attention is directed toward cues predictive of reward, and unrewarded or irrelevant signals are ignored (Gottlieb, 2012; Mackintosh, 1975). Such filtering reduces representational and learning costs, though it may limit sensitivity to rewarding opportunities (Kolling et al., 2016).

Selective behavioral strategies are associated with distinct frontal representations, characterized by strong responses to reward-associated cues and diminished responses to non-rewarded stimuli (Ebitz & Hayden, 2016). Within this framework, frontal cortex integrates sensory, motivational, and feedback signals to support the selection and maintenance of efficient behavioral policies (Botvinick et al., 2001; Chen et al., 2024; Kolling et al., 2016; Rushworth et al., 2011). By preferentially amplifying predictive inputs and down-weighting irrelevant ones, frontal circuits bias behavior toward reliable cues and stable response strategies (Dayan et al., 2000; Niv et al., 2015; Shenhav et al., 2016; Kim et al., 2021; Narayanan et al., 2006; Peters et al., 2022; Sridharan et al., 2008; Totah et al., 2009).

The frontal cortex, and in particular the anterior cingulate cortex (ACC), is a principal target of the claustrum, positioning the claustrum as a candidate modulator of frontal representations and control policies (Atlan et al., 2016, 2018a; Wang et al., 2023; Zingg et al., 2018). Through recruitment of cortical interneurons, claustral activity modulates cortical states and large-scale coordination, influencing synchronization and slow oscillatory dynamics linked to arousal, memory, and network organization (Atlan et al., 2024; Do et al., 2024; Marriott et al., 2024; Jackson et al., 2018; Mcbride et al., 2023; Narikiyo et al., 2020; Zahacy et al., 2024). Importantly, the impact of claustral output on cortex is strongly state dependent, varying with arousal and the phase of ongoing cortical activity (Marriott et al., 2024; Narikiyo et al., 2020).

Consistent with this role, the claustrum has been implicated in motor planning, response inhibition and cognitive control under demanding conditions (Atlan et al., 2018a; Chevée et al., 2022; Liu et al., 2019a; White et al., 2020). In particular, projections from the claustrum to the ACC support response inhibition and adaptive control (Atlan et al., 2024), and disruption of claustrum activity impairs the formation of medial prefrontal assemblies during attentional set-shifting (Fodoulian et al., 2020). Together, these findings raise the possibility that claustral input contributes to organizing frontal dynamics required for adaptive control.

Despite these advances, a key question remains unresolved: does the claustrum primarily support moment-to-moment behavioral control, such as salience detection or response inhibition, or does it play a broader role in shaping and stabilizing frontal control strategies across experience? While early hypotheses emphasized transient encoding of salient events (Crick & Koch, 2005; Smythies et al., 2014), accumulating evidence indicates that claustral influence extends beyond immediate control to the coordination of cortical and behavioral states in a context-dependent manner (Atlan et al., 2024; Lamsam et al., 2024; Marriott et al., 2024; Mcbride et al., 2023; Narikiyo et al., 2020; Zahacy et al., 2024). Notably, sensory representations in claustral projections to the ACC correlate with inter-individual differences in behavioral policy (Atlan et al., 2024), raising the possibility that claustral activity contributes to the consolidation of selective control strategies rather than their online execution.

We recently showed that transitioning from autonomous training to head-fixed task execution induces a reorganization of behavioral response profiles, reflecting the need to stabilize efficient strategies under altered temporal and motivational constraints (Peretz-Rivlin et al., 2026). This transition therefore provides a natural opportunity to test whether claustral input instructs the consolidation of frontal representations that support selective behavioral policies. To address this question, we combined optogenetic perturbation of ACC-projecting claustral neurons with large-scale Neuropixels recordings from frontal cortex during head-fixed task performance after autonomous training, enabling us to assess how disrupting claustral input affects behavioral selectivity and the organization of frontal population dynamics.

## Results

### Behavioral task and response patterns in control mice

To examine how claustral input relates to behavioral selectivity and frontal cortical encoding during task performance, we established an experimental framework combining large-scale Neuropixels recordings from frontal cortex with optogenetic perturbation of anterior cingulate–projecting claustral neurons during performance in the ENGAGE task (Atlan et al., 2024; Figure 1A). Mice were first extensively trained in automated home-cage environments and subsequently transitioned to a head-fixed configuration that enabled simultaneous behavioral monitoring, neural recording, and optogenetic stimulation (Figure 1B).

**Figure 1.**
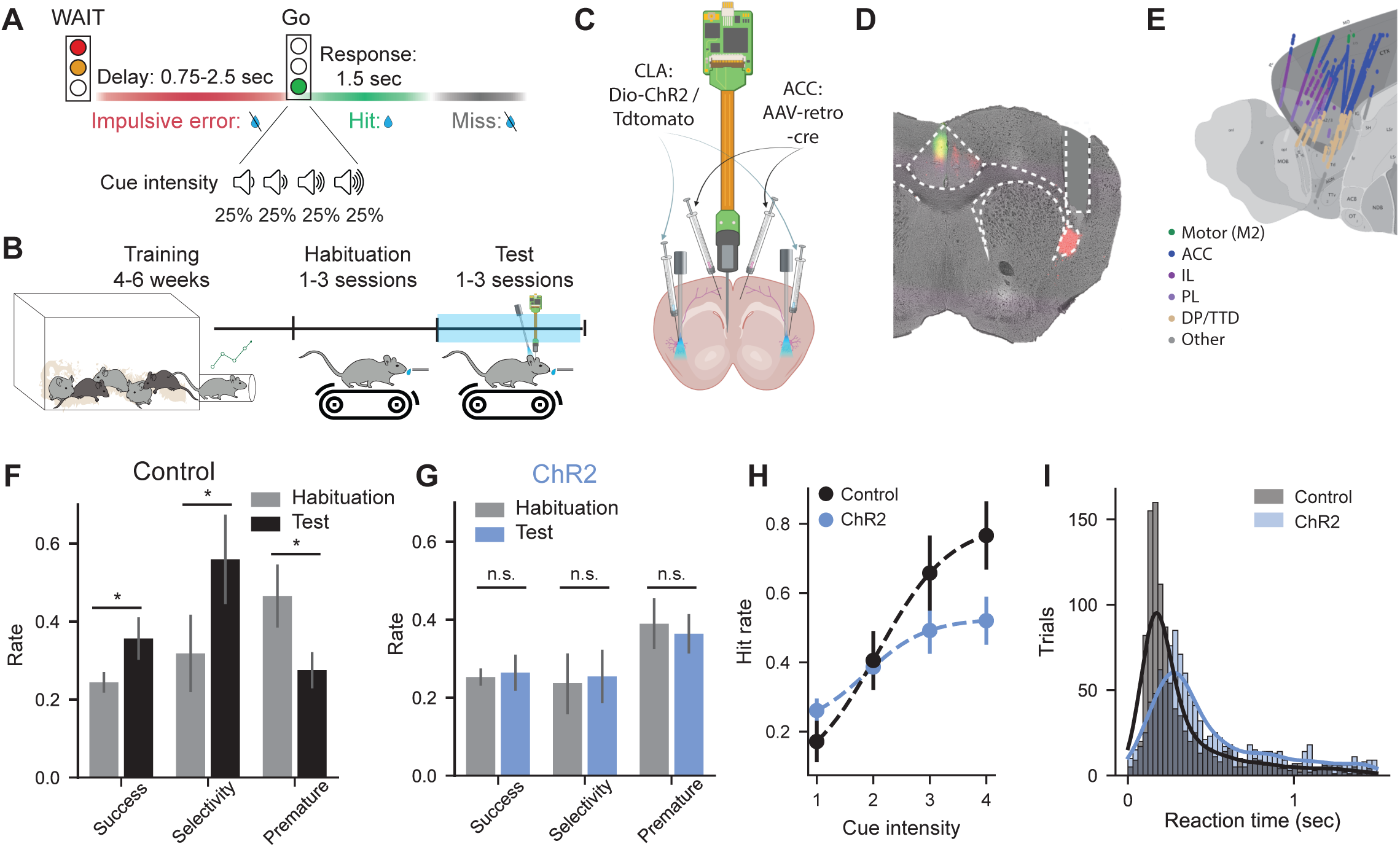
Experimental design, neural interface, and behavioral performance in the ENGAGE task. (**A**) Schematic of the ENGAGE task trial structure. Trials initiate with a WAIT signal, followed by a variable delay (0.7–2.5 s). The GO cue is presented at four logarithmically spaced intensities, embedded in a masking tone cloud in 50% of trials. Licking within the 1.5-second response window triggers reward. (**B**) Experimental timeline. Mice underwent automated training in the home cage (4–6 weeks) to acquire task contingencies before transfer to the head-fixed setup. The protocol included 1–3 habitua-tion sessions followed by 1–3 test sessions combining behavioral testing with optogenetics and Neuropixels recording. (**C**) Viral strategy and recording configuration. AAV-retro-Cre was injected into the Anterior Cingulate Cortex (ACC), and Cre-de-pendent opsins (AAV-DIO-ChR2) or fluorophores (AAV-DIO-tdTomato) were injected bilaterally into the claustrum. Optic fibers were implanted above the claustrum, and Neuropixels probes were acutely inserted into the ACC. (**D**) Representative histology showing viral expression and fiber placement in the claustrum, alongside a DiO-labeled Neuropixels track in the ACC. (**E**) Reconstructed trajectories of Neuropixels probe insertions across all recordings, color-coded by anatomical region. (**F**) Comparison of behavioral metrics (Success rate (p=0.016, linear mixed effect model), Premature rate (p=0.023, linear mixed effect model), Selectivity Index (p=0.018, linear mixed effect model) in control mice between habituation (n=8 session) and test sessions (n=10 sessions). Data presented as mean ± SEM. (**G**) As in (F), for the ChR2 group (Habitua-tion: n=10 sessions; Test: n=14 sessions, all comparisons are n.s. in linear mixed effect models). (**H**) Psychometric curves comparing Control (n=10 sessions) and ChR2 (n=14 sessions) groups during test sessions. Hit rates are plotted as a function of sound intensity (excluding premature trials). Data represent mean ± SEM. Interaction terms were significant at higher cue levels (Linear mixed effect model ‘Hit∼intensity*group + 1/subject’; intensity 3: β = 0.26 ± 0.09, z = 2.71, p = 0.007; intensity 4: β = 0.34 ± 0.09, z = 3.56, p < 0.001) (**I**) Reaction time distributions for correct (hit) trials in test sessions, pooled across all animals and trials for Control vs. ChR2 groups (Mann–Whitney U test, U = 3.77 × 10⁵, p = 2.6 × 10^-^⁴⁵).

Following transfer to head fixation, task contingencies were unchanged, but trials were externally paced and limited in number, requiring animals to express previously acquired task rules under altered temporal and motivational constraints. This transition is associated with the stabilization of a response strategy characterized by selective responding to high-intensity GO cues and reduced premature actions (Peretz-Rivlin et al., 2026; Figure S1A-C). This selective strategy supports efficient performance under constrained trial availability, defining a behavioral regime in which cortical encoding and the impact of claustral perturbation can be directly assessed.

To probe the impact of claustral disruption on cortical encoding and strategy consolidation, we implemented high-density recordings of frontal activity together with optogenetic stimulation of claustral projection neurons (Figure 1C). Neuropixels recordings were obtained from frontal cortical regions, primarily the anterior cingulate cortex (ACC; Figures 1C-E). In parallel, ACC-projecting claustral neurons (ACCp) were optogenetically stimulated during defined epochs of the task. Optogenetic stimulation was interleaved either around the *WAIT* signal (25% of trials) or the period surrounding the *GO*-cue (25%), corresponding to task phases associated with response inhibition and motor preparation, respectively (Figure S1D). This design allowed us to test whether disrupting the timing of claustral input during defined task epochs interferes with the consolidation of selective behavioral strategies and their associated frontal cortical representations.

### Temporally misaligned claustral signaling interferes with the consolidation of selective strategies

Group differences emerged during the test sessions in which optogenetic stimulation was applied. While control mice exhibited higher success rates, greater cue selectivity and improved response efficiency during test sessions relative to habituation (Figure 1F; Figure S1A-C), experimental mice expressing ChR2 in ACC-projecting claustral neurons failed to show comparable improvements in these measures (Figure 1G).

Psychometric analyses of performance during test sessions reinforced this distinction. ChR2 mice exhibited flatter intensity-response functions, indicating reduced sensitivity to *GO* cue intensity (Figure 1H). Control animals also responded with shorter latencies than ChR2 mice, consistent with more efficient sensory-to-motor transformations (Figure 1I). Together, these results demonstrate that perturbation of claustrum output interferes with the consolidation of efficient, selective task strategies across sessions, resulting in persistently elevated impulsive errors and reduced performance stability.

Notably, claustrum perturbation timed to defined task epochs did not drive detectable trial-by-trial effects on behavior (Figure S1F). Across both control and ChR2 cohorts, stimulated and unstimulated trials did not differ in success rate, premature responding, omissions, or cue selectivity. Thus, although perturbation of ACC-projecting claustrum neurons disrupted the consolidation of a selective task strategy across sessions, it did not acutely bias online behavioral execution. This dissociation was observed despite robust physiological engagement of the claustro-frontal circuit. Validation recordings confirmed that optogenetic stimulation of ACCp neurons reliably modulated cortical activity, with the majority of responsive units exhibiting significant downregulation (42%), while a smaller proportion was upregulated (24%) (Figures S1G-I), consistent with previous reports (de la Torre-Martínez et al., 2025).

### Sensory and Motor Representations in the ACC

Having observed that claustral perturbation disrupted the consolidation of selective behavioral strategies, we next characterized the encoding of cues and actions by frontal cortical populations, in order to establish a reference for interpreting the effects of claustral perturbation.

We first characterized task-related single-unit responses in control mice using large-scale Neuropixels recordings from frontal cortex. Rastermap-based ordering (Stringer et al., 2025) of peri-stimulus time histograms (PSTHs) revealed 19 reproducible functional clusters, which were grouped using hierarchical clustering into sensory, premotor, reward-related, and task non-responsive (task NR) populations based on their temporal alignment to task events (Figures 2A, B, S2A, S2B). To ensure valid comparison between groups, clustering was performed on neural activity pooled from control and ChR2 mice, and only trials without laser stimulation were included to isolate task-related cortical activity.

**Figure 2.**
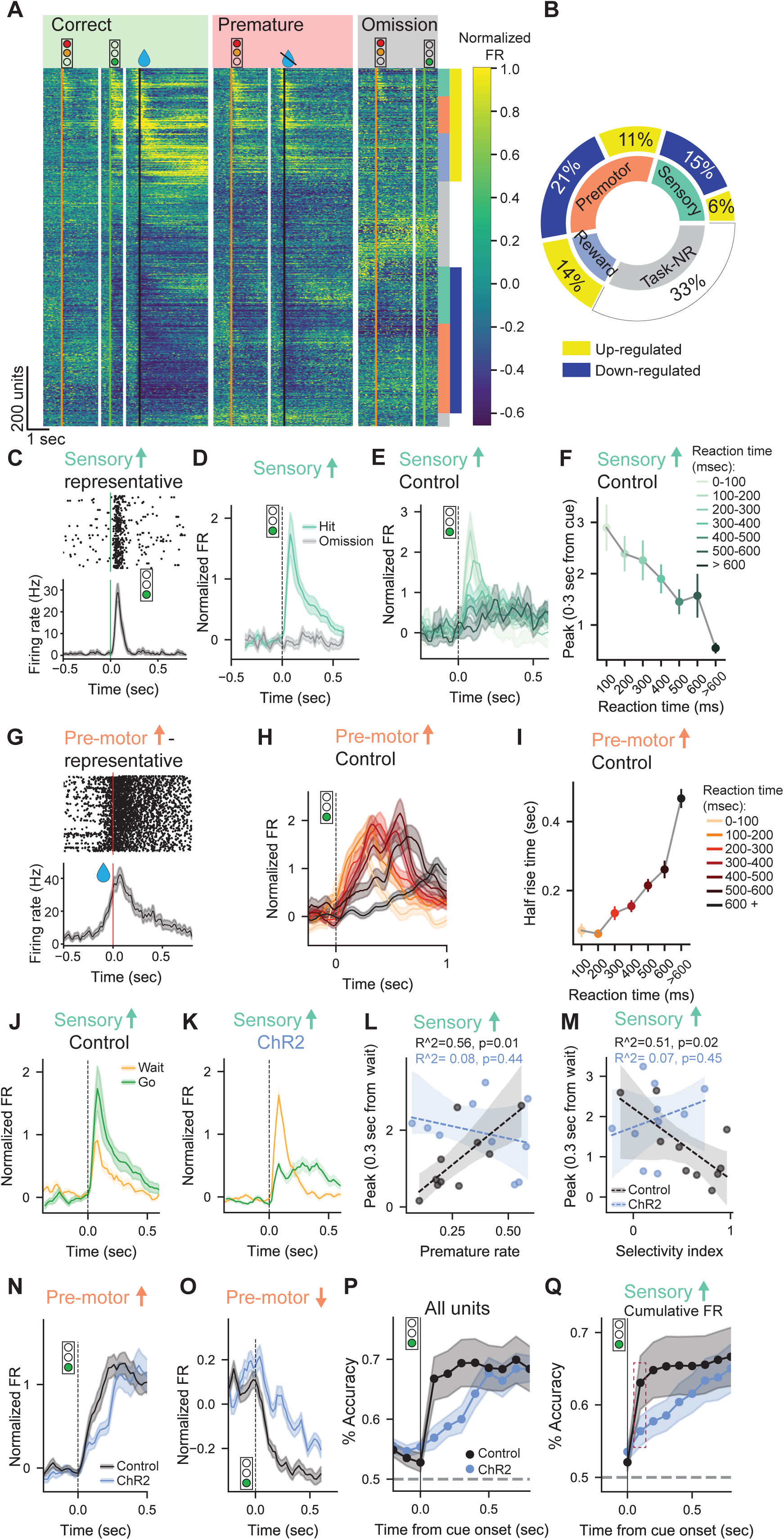
Claustral perturbation disrupts sensory-to-motor transformations in frontal cortex. (**A**) Heatmap showing the normalized firing rate of recorded units from control mice (n=1846 units) averaged across relevant task events. For correct trials, aligned events include the WAIT signal, GO-cue, and rewarded lick. For premature trials, alignment includes the WAIT signal and unrewarded lick. For omission trials, responses to the WAIT signal and GO-cue are shown. Units were sorted using the ‘Rastermap’ algorithm (Stringer et al., 2025) and color-coded by functional classification: sensory, premo-tor, reward, or other (classification defined in Supplementary Figure 2). (**B**) Pie chart showing the distribution of recorded units into categories, further divided into upregulated and downregulated subtypes (see methods). (**C**) Raster plot and peristimulus time histogram (PSTH) aligned to the GO-cue in hit trials for an example unit classified as sensory-upregulat-ed. (**D**) Peristimulus time histogram (PSTH) of sensory upregulated units from Control mice (n=66 units) aligned to GO-cue, by outcome (Hit vs. Omission) (**E**) Average response of sensory-upregulated units from controls (n = 66 units) to the GO-cue in hit trials, separated by reaction time quantiles. Lines represent means; shaded areas indicate SEM. (**F**) Peak normalized firing rate in the first 300 ms following GO-cue onset in hit trials, plotted against reaction time bins. Points repre-sent mean values; error bars indicate SEM (n = 66 units units). (**G**) Raster plots and PSTHs aligned to the GO-cue in hit trials for a representative unit classified as premotor. (**H**) Average response of all premotor-upregulated units from control mice (n = 115 units) to the GO-cue in hit trials, separated by reaction time quantiles. (**I**) Half-rise time of premotor-upregulat-ed unit responses following GO-cue onset in hit trials, plotted against reaction time bins. Points represent means; error bars indicate SEM (n = 115 units). (**J-K**) Comparison of responses to the WAIT signal versus the GO-cue in sensory-upregulated units: (**J**) Controls (n = 66 units) and (**K**) ChR2 (n = 127 units). Lines show means; shaded areas, SEM. (**L-M**) Session-wise (control n=10, ChR2 n=14 sessions) correlations between peak sensory-upregulated responses to the WAIT signal (0–300 ms) and (**L**) premature error rate or (**M**) selectivity index. (**N-O**) Average responses aligned to the GO-cue in hit trials, comparison of Controls and ChR2, for: (**N**) premotor-upregulated units (129 vs. 115 units), and (**O**) premotor-downregulated units (238 vs. 294 units). Lines show means; shaded areas, SEM. (**P-Q**) Decoding performance for predicting hit vs. omission trials following GO-cues from: (**P**) full-population firing rates, (Q) Cumulative firing rates of sensory-upregulated units (8% of recorded neurons). (Control n=10sessions, ChR2 n=14 sessions).Red dashed rectangular mark the time point used in Figure S3E.

In control mice, a prominent population of sensory units exhibited rapid, transient responses aligned to *GO*-cue onset (Figure 2C). These responses were action-contingent, robust during hit trials but absent during omissions, indicating encoding of decision-relevant sensory evidence rather than passive sensation (Figure 2D). Sensory response profiles scaled systematically with reaction time: fast responses were associated with sharp, high-amplitude peaks, whereas slower responses showed broader, lower-amplitude activity (Figures 2E, F). Notably, cumulative sensory activity remained constant across trials, consistent with an evidence-accumulation–like process linking sensory encoding to action timing (Figure S3A) (Shadlen & Newsome, 2001).

Downstream premotor units displayed ramping activity preceding lick onset (Figure 2G). When aligned to the *GO*-cue, ramp duration scaled linearly with reaction time (Figures 2H, I), whereas alignment to lick onset revealed stereotyped, time-invariant profiles (Figure S3B). Together, these dynamics delineate a canonical sensory-to-motor sequence in frontal cortex, whereby accumulated sensory evidence recruits premotor ensembles that ramp to an action threshold (Hanks et al., 2014).

Consistent with the absence of within-session behavioral effects, claustrum stimulation did not measurably alter task-evoked neural responses on stimulated trials (Figures S3C-H). Nevertheless, clear group-level differences were observed in behavior (Figure 1) as well as the corresponding neural representations between control and ChR2 mice. Sensory units in ChR2 mice showed attenuated responses to the *GO*-cue and exaggerated responses to the *WAIT*-signal (Figures 2J, K; S3I, J), indicating abnormal prioritization of non-instructive cues at the expense of task-relevant sensory evidence. In control mice, stronger *WAIT*-signal encoding was associated with increased impulsivity and reduced behavioral selectivity, whereas these relationships were absent in ChR2 mice (Figures 2L, M), suggesting a breakdown in the linkage between sensory representations and behavioral policy.

Consistent with impaired sensory–motor coupling, premotor recruitment was delayed in ChR2 mice (Figures 2N, O), paralleling their prolonged reaction times. Logistic regression decoding further demonstrated that trial outcomes could be predicted rapidly from frontal population activity in control mice but only gradually in ChR2 mice (Figure 2P). Restricting decoding to sensory-responsive units reproduced this group difference despite using a small fraction of recorded neurons, indicating that degraded sensory encoding alone could account for delayed outcome predictability (Figure 2Q). Across sessions, early decoding accuracy from sensory units correlated with behavioral selectivity, directly linking rapid and efficient frontal sensory encoding to the stabilization of efficient strategies (Figure S3K).

Together, these results show that claustral perturbation degrades task-relevant sensory encoding, exaggerates responses to non-instructive cues, and delays premotor recruitment, disrupting the transformation of frontal representations into stable behavioral strategies.

### ACC Background Activity Gates Impulsive and Omissive Behavior

Beyond phasic sensory–motor transformations, effective behavioral control in decision making tasks also depends on the tonic baseline state of frontal cortex prior to cue presentation. The ACC regulates sensory–motor flow in part through task-agnostic neuronal activity that is not selectively tuned to specific task events (Kim et al., 2021).

In control mice, pre-trial firing rates of this ‘Task-NR’ population were predictive of subsequent behavioral outcomes: low baseline activity preceded premature responses, high baseline activity preceded omissions, and intermediate activity preceded correct trials (Figure 3A). In ChR2 mice, elevated baseline firing before omission trials persisted, but the distinction between baseline activity preceding correct and premature trials was lost (Figure 3B). This loss of separation resulted primarily from reduced baseline firing during correct trials in ChR2 mice (Figure 3C).

**Figure 3.**
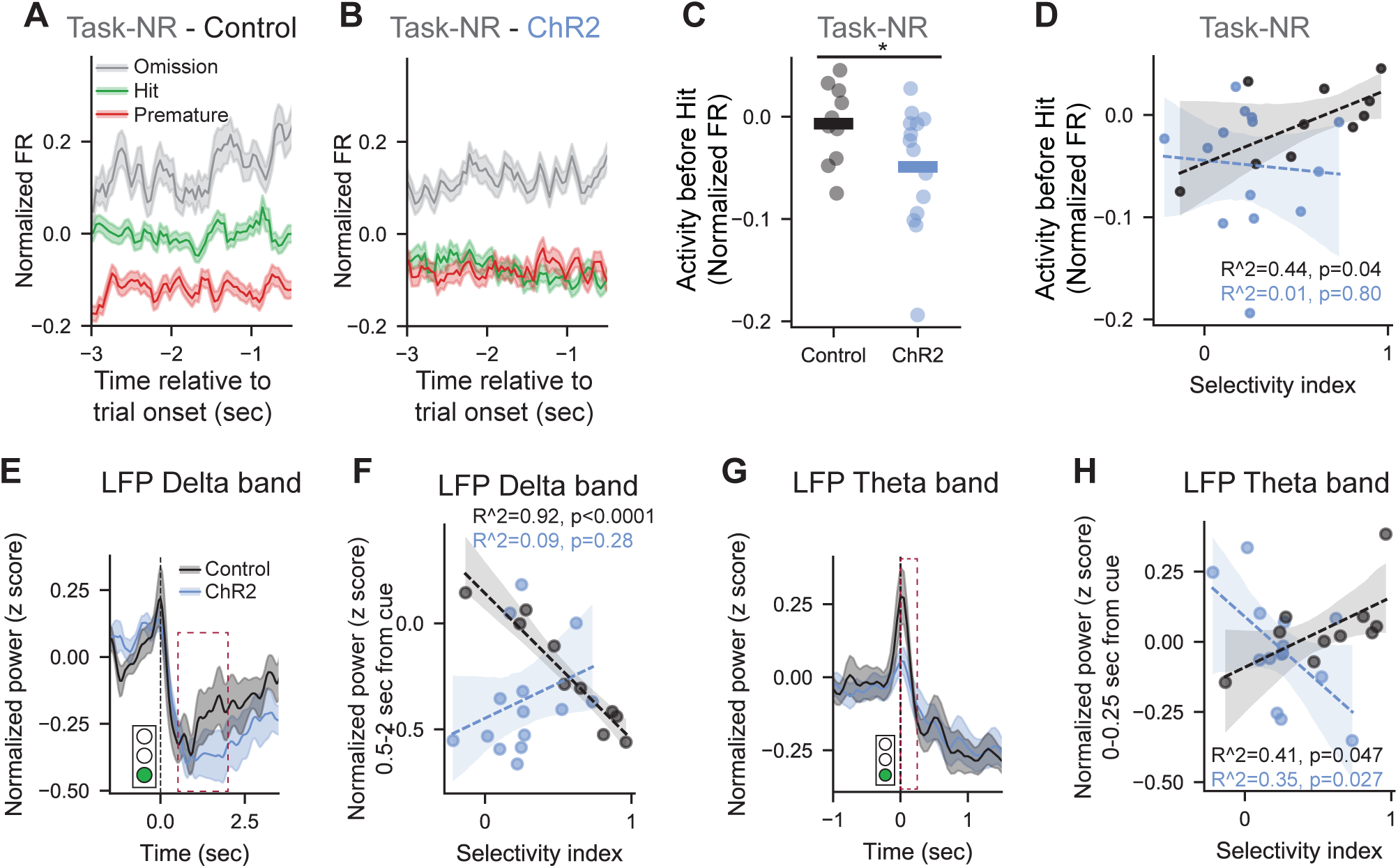
Claustral perturbation disrupts the coupling between frontal background activity, LFP dynamics, and behavioral selectivity. (**A**) Average normalized firing rate during the pre-trial baseline period for ‘Task-NR’ units in Control mice (n = 370 units), separated by subsequent trial outcome. (**B**) Same as (A) for ChR2 mice (n = 340 units). (**C**) Mean baseline activity of the ‘Task-NR’ population in hit trials, by group. Each point represents one session (Controls: n = 370 units, 10 sessions; ChR2: n = 340 units, 14 sessions). Welch t-test: t = 2.106, p = 0.0469. (**D**) Baseline activity in hit trials plotted against behavioral selectivity index. Each point represents one session; separate regression lines were fit to each group of mice. Control: R² = 0.44, p = 0.04; DIO-ChR2: R² = 0.01, p = 0.80. (**E-F**) Delta-band (0.5–4 Hz) power following GO-cue onset in hit trials. (**E**) Time course of changes in normalized power. Red dashed rectangular mark the time points averaged in (F). (**F**) Mean power (0.5–2 s) versus selectivity index, showing strong correlation in controls (R² = 0.92, p < 0.0001) but not ChR2 mice. (**G-H**) Theta-band (6–11 Hz) power following GO-cue onset in hit trials. (**G**) Time course of changes in normalized power. Red dashed rectangular mark the time points averaged in (H). (**H**) Mean power (0–0.25 s) versus selectivity index, correlated in both groups (controls: R² = 0.41, p = 0.047; ChR2: R² = 0.35, p = 0.027).

In control mice, but not in ChR2 mice, pre-trial activity of the ‘Task-NR’ population preceding correct responses correlated positively with the behavioral selectivity index, indicating that animals adopting more selective strategies also maintained higher baseline activity before cue presentation, potentially supporting stronger inhibitory control (Figure 3D). Notably, the dynamics of this population mirror those previously observed in ACC-projecting claustral neurons (Atlan et al., 2024).

Together, these results indicate that in mice whose claustral activity is intact, tonic background activity in this task-agnostic population gates sensory–motor transformation, consistent with prior work (Kim et al., 2021). High baseline activity likely suppresses motor output, leading to omissions, whereas intermediate activity permits appropriate responding without impulsive actions. Disruption of this modulatory structure in ChR2 mice therefore reveals an additional deficit in inhibitory gating that undermines adaptive behavioral control.

### Delta and Theta Band LFP Dynamics Reflect Behavioral Selectivity

Building on the observed disruption of sensory-motor coupling by claustral perturbation, we next asked whether local field potential (LFP) dynamics also reflected this deficit. Frontal theta (6-11 Hz) and delta (0.5–4 Hz) oscillations have been strongly linked to cognitive control, motor preparation and inhibitory gating (Cavanagh & Frank, 2014; Harmony, 2013). Theta power typically increases with cue-induced readiness, whereas transient delta suppression accompanies release from inhibition and efficient action initiation (Cavanagh & Frank, 2014; Cavanagh & Shackman, 2015; Harmony, 2013; Igarashi et al., 2013; Nigbur et al., 2011; Robble et al., 2021; Wessel et al., 2020).

In control mice, correct *GO-*cue responses were associated with a transient increase in frontal theta power at cue onset, an effect that was absent in ChR2 mice (Figure 3E). Theta magnitude correlated positively with behavioral selectivity in controls, whereas in ChR2 mice this relationship was inverted, consistent with the deployment of a less effective behavioral strategy by experimental mice (Figure 3F). In both groups, delta power decreased following *GO*-cues that elicited correct responses (Figure 3G), reflecting the release from inhibition. However, only in control mice did the magnitude of delta suppression scale with behavioral selectivity, with stronger suppression accompanying more efficient strategies that balanced inhibition and action (Figure 3H).

Together, these results show that frontal theta-delta dynamics track behavioral selectivity, reflecting coordinated cortical state transitions during task performance. Disruption of these oscillatory signatures in ChR2 mice indicates that claustral input is required to organize large-scale cortical rhythms that support efficient and selective behavioral control.

### Claustrum Perturbation Induces Global Shifts in Cortical Population States during response inhibition

To determine whether claustral perturbation alters the global organization of cortical population activity, we applied a supervised dimensionality-reduction approach to neural activity across all task epochs. Population activity was segmented into discrete task epochs and embedded into a low-dimensional latent space using CEBRA (Schneider et al., 2023), enabling an unbiased comparison of task-related population dynamics between control and ChR2 mice (Figure 4A).

**Figure 4.**
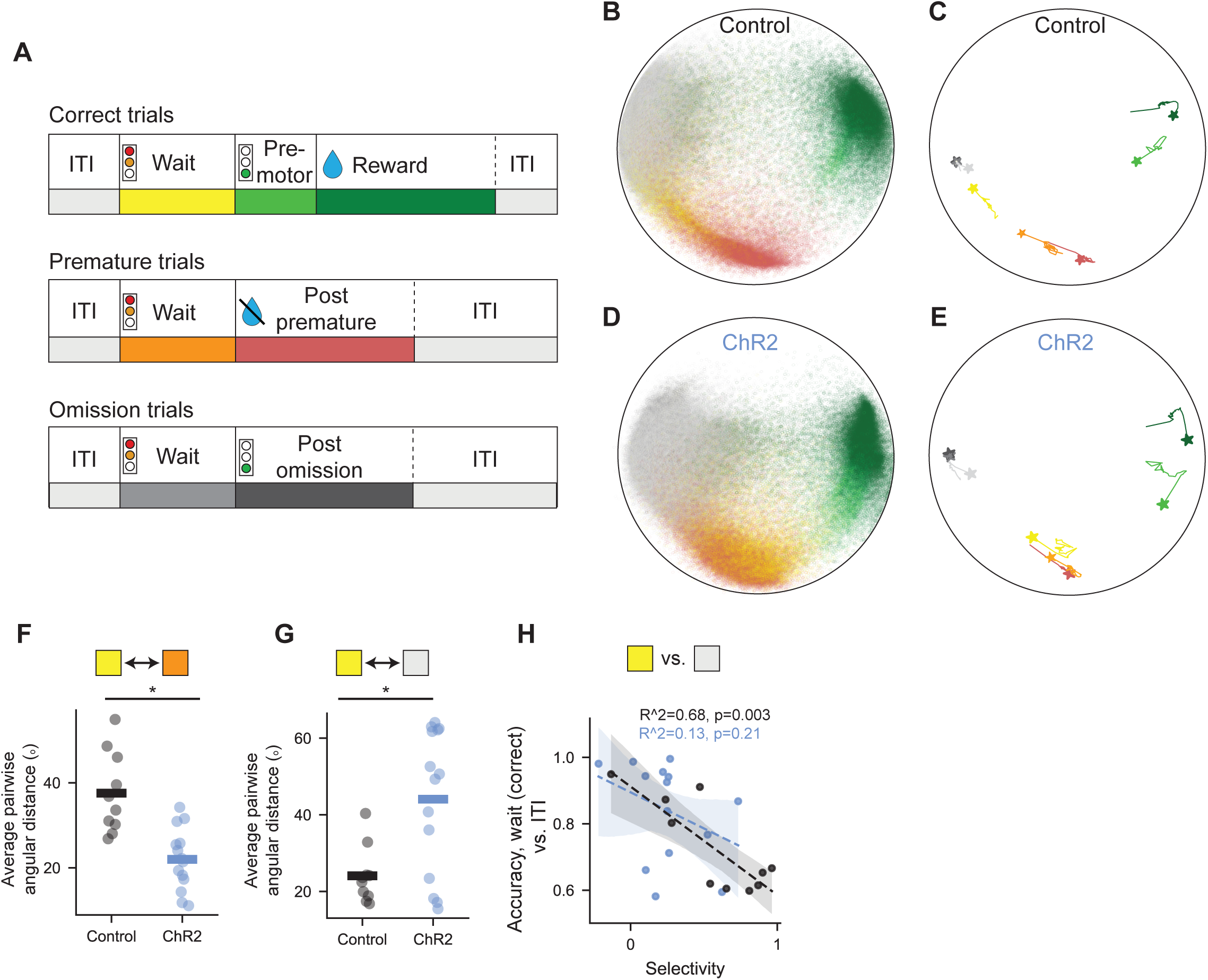
Claustral perturbation alters global population states during response inhibition. (**A**) Task epoch labels assigned to each 50-ms time point in the experiment. Only time points without laser stimulation were taken into the analysis.(**B**) Three-dimensional latent embedding of all time points from control sessions, computed using the supervised CEBRA algorithm (Schneider et al., 2023) and color-coded by task epoch (legend in A). (**C**) Mean trajectories for each labeled epoch from (B); asterisks indicate phase initiation. Colors correspond to (A). (**D–E**) Same as (B–C) for ChR2 sessions. (**F**) Mean pairwise Angular distance between embedded points from the inhibitory period (after the WAIT signal) in correct and in premature trials, by group (Welch t-test: t = 4.352, p = 0.0005). (**G**) Mean pairwise angular distance between embedded points from the inhibitory period in correct trials and inter-trial interval (ITI), by group (Welch t-test: t = -3.611, p = 0.002). (**H**) Accuracy of a k-nearest neighbor (KNN) decoder distinguishing the inhibitory period in correct trials from ITI epochs plotted against the behavioral selectivity index. Each point represents a session; separate regression lines are shown for each group (Controls: R² = 0.68, p = 0.003; ChR2: R² = 0.13, p = 0.21).

This global embedding revealed a robust organization of population activity by task epoch in both groups, with similar overall geometry of sensory, motor, and outcome-related states (Figures 4B–E; see supplementary data for interactive embedding), indicating that claustral perturbation does not broadly disrupt task-related population structure. Within this preserved global organization, however, a specific divergence emerged in the representation of the inhibitory period between the *WAIT* signal and the *GO*-cue, a time-point during which mice were required to withhold responding.

In control mice, neural activity during the inhibitory period clustered near unresponsive states, such as the inter-trial interval (ITI) and omission epochs (Figures 4B, C), consistent with a disengaged or inhibited cortical state. In contrast, in ChR2 mice, inhibitory-period activity was shifted toward representations associated with premature errors and impulsive responses (Figures 4D, E). This shift indicates premature recruitment of motor-related dynamics even on trials in which mice successfully withheld responding, providing a mechanistic explanation for the elevated impulsivity observed behaviorally (Figure 1J). Notably, this population-level bias is consistent with the exaggerated WAIT-signal encoding, impaired tonic inhibitory gating, and delayed sensory–motor coupling observed at the single-unit and oscillatory levels (Figures 2 and 3).

We quantified these group differences by computing the angular distance in latent space between inhibitory-period activity in correct trials and activity during premature or ITI epochs for each recording session. In control mice, inhibitory-period activity was more distant from premature-error representations (Figure 4F), whereas in ChR2 mice it was more distant from ITI representations (Figure 4G). Thus, in control animals, the inhibitory period more closely resembled an unresponsive state, while in ChR2 mice it was already biased toward action-related representations.

To further assess the behavioral relevance of these representations, we trained a k-nearest neighbor classifier to distinguish between task epochs based on embedded population activity. In control mice, classifier accuracy for distinguishing ITI from inhibitory-period activity was negatively correlated with behavioral selectivity: animals exhibiting higher selectivity showed greater overlap between these states. This relationship was absent in ChR2 mice (Figure 4H), indicating a breakdown in the coupling between cortical state organization and behavioral strategy.

Together, these results show that control mice, particularly those with high behavioral selectivity, maintain a disengaged cortical state during the inhibitory period, consistent with effective response inhibition and a selective suppression of non-instructive cues. In contrast, claustral perturbation disrupts the establishment of this state, biasing cortical dynamics toward action-related representations during periods requiring inhibition and thereby reducing behavioral efficiency.

## Discussion

In this study, we identify a central role for claustral projections to the anterior cingulate cortex (ACC) in enabling the stabilization of adaptive behavioral strategies under demanding task conditions. When claustro–ACC signaling was perturbed, mice failed to converge on the efficient response policy expressed by control animals, instead showing persistently elevated premature responses and reduced cue selectivity during head-fixed performance (Figures 1F–I). This divergence reflects a disruption in the control processes that determine whether actions are withheld or released, pointing to a failure of response inhibition.

Functions in adaptive behavioral control have been attributed to frontal cortex, basal ganglia, thalamic nuclei, and neuromodulatory systems, each of which has been implicated in decision making, response inhibition, or reinforcement-driven learning. These contributions, however, have largely been characterized in terms of online control variables such as action selection, value updating, or moment-to-moment gating of motor output, or through broad modulation of cortical excitability. In contrast, our findings identify a distinct role for the claustrum in stabilizing frontal population dynamics and control states across experience. Disrupting claustro–ACC signaling did not abolish task execution, sensory–motor transformations, or trial-by-trial performance, but selectively interfered with convergence onto an efficient, selective behavioral strategy. These results suggest that the claustrum does not directly implement control policies or gate individual actions, but instead acts as a circuit element that maintains the conditions under which frontal control strategies can consolidate and stabilize over experience.

At the neural level, impaired strategy stabilization was accompanied by a coordinated set of alterations in frontal processing. Claustral perturbation distorted the relative weighting of instructional sensory signals, exaggerating responses to the WAIT cue while attenuating GO-evoked activity (Figures 2J-M), destabilized population states during periods requiring response withholding, and weakened the coupling between cortical dynamics and behavioral outcomes (Figures 3; 4). These changes occurred despite preserved sensory-evoked and motor-related responses and intact trial-level performance (Figure S1F), indicating that claustro–ACC input is required to organize and maintain frontal control states that support selective, inhibition-based behavior, rather than directly driving sensory processing or motor execution.

Behavioral specificity was further supported by the preservation of core task capacities in claustrum-perturbed mice. Experimental animals retained accurate responses to high-intensity cues (Figure 1H), exhibited intact sensory-evoked and motor-related activity on individual trials (Figures 2J-K, 2N-O), and showed no evidence of impaired knowledge of task contingencies (Berditchevskaia et al., 2016; Kim et al., 2021). Instead, deficits emerged selectively in contexts requiring sustained suppression of premature responding. Control mice established a performance policy that was based on maintaining a strong inhibitory gate through the WAIT period and releasing action reliably only once sufficient sensory evidence accumulated, whereas ChR2-expressing mice exhibited elevated false alarms and reduced cue selectivity (Figures 1F-I).

At the neural population level, this phenotype was associated with destabilization of frontal population states normally linked to response inhibition. Task-agnostic (“task-NR”) neurons failed to segregate in activity prior to correct and premature actions (Figures 3A-D), consistent with disrupted inhibitory gating implemented through background frontal activity (Kim et al., 2021). Low-frequency theta–delta dynamics, previously linked to frontal control signaling and inhibitory suppression (Cavanagh & Frank, 2014; Harmony, 2013), became decoupled from behavioral outcomes (Figures 3E-H), and population activity during the WAIT period was biased toward action-related states (Figures 4B-G). Importantly, these effects occurred within an otherwise preserved global organization of task-related population activity, indicating a selective disruption of control-related dynamics, rather than a collapse of task representations.

Within this control regime, selective responding was supported by a coordinated relationship between cue encoding and population state. In control mice, encoding of the *WAIT* signal was relatively weak and frontal population activity during the *WAIT* period remained in a non-action state continuous with the inter-trial interval, maintaining suppression of motor-related dynamics until sufficient sensory evidence was available. Robust and temporally precise responses to the *GO* cue reliably drove transitions from inhibition to action. In contrast, claustral perturbation profoundly altered this relationship: ChR2-expressing mice exhibited exaggerated encoding of the *WAIT* signal alongside attenuated *GO*-cue responses (Figures 2J–M), accompanied by a shift of population dynamics toward an action-related regime during the waiting period (Figures 4D–E). Rather than sustaining a non-action state, strong *WAIT* encoding in the absence of appropriate claustral input was associated with premature engagement of motor-associated dynamics, undermining the network conditions required for effective response inhibition (Miller & Cohen, 2001; Akam et al., 2021; Totah et al., 2009).

Critically, these effects did not manifest as acute, trial-by-trial biases in behavior. Despite robust modulation of frontal firing rates, optogenetic stimulation did not reliably alter performance on stimulated trials (Figure S1F). Instead, its impact emerged across sessions, interfering with the consolidation of efficient, selective strategies. This dissociation argues against a purely online gating or salience-detection role for the claustrum (Crick & Koch, 2005; Smith et al., 2019), and instead supports a model in which claustral input contributes to the stabilization of frontal population dynamics over experience (Atlan et al., 2024; Terem et al., 2020).

Together with prior work showing that claustral disruption preferentially impairs performance under high cognitive demand while sparing simpler, previously consolidated behaviors (Atlan et al., 2018b; Liu et al., 2019b; Terem et al., 2020; White et al., 2020), these findings point to a general mechanistic role for the claustrum as a coordinator of cortical control states. Rather than encoding a specific cognitive variable such as attention, salience, or inhibition, the claustrum acts as a hub that aligns frontal population dynamics with behavioral demands (Jackson et al., 2018; Mcbride et al., 2023; Zahacy et al., 2024). When this coordination is intact, frontal cortex consolidates representations that are stable, selective, and predictive of behavior; when disrupted, frontal cortex retains overall encoding of task structure, and generates actions, but fails to stabilize new control regimes required for efficient, selective behavior upon changes to task context and behavioral constraints (Brockett et al., 2020; Fodoulian et al., 2020).

Several directions for future work follow naturally from these findings. Because neural recordings were obtained only after transfer to head fixation, the present study does not directly capture how frontal control states emerge during initial training and habituation. Longitudinal recordings spanning the full learning trajectory will be necessary to determine whether claustral input is required primarily for the acquisition, refinement, or maintenance of inhibitory control regimes. In addition, complementary loss-of-function approaches, including temporally precise inhibition of ACC-projecting claustral neurons, will be important for testing whether suppression of claustral input produces comparable disruptions in frontal state stabilization.

In summary, this work identifies claustral projections to the ACC as a critical circuit for stabilizing frontal population dynamics that support response inhibition and selective behavior. By integrating large-scale neural recordings with targeted circuit perturbation, we show that claustrum–ACC interactions support the neural conditions under which efficient control strategies can be consolidated across experience. These findings position the claustrum as a subcortical scaffold for adaptive control and provide a mechanistic framework for understanding how its dysfunction may contribute to impulsivity and impaired cognitive flexibility.

## Methods

### Animals

All mice described in this study were male C57BL/6JOLAHSD obtained from Harlan Laboratories, Jerusalem, Israel. Mice were housed in groups of littermates and kept in a SPF (specific pathogen-free) animal facility under standard environmental conditions: temperature (20–22 °C), humidity (55 ± 10%), and 12–12 h light/dark cycle, with ad libitum access to water and food. Mice were randomly assigned to experimental groups. All experimental procedures, handling, surgeries, and care of laboratory animals used in this study were approved by the Hebrew University Institutional Animal Care and Use Committee (NS-20-16301-4). A total of 14 mice were injected and tested. Of these, 7 received AAV-DIO-ChR2 and 5 received AAV-DIO-TdTomato, and were included in the analysis. Two mice were excluded due to off-target viral expression outside the claustrum, as confirmed by post-experimental histological verification.

### Preparatory stereotactic surgery for viral injections and cannula implantation

Induction and maintenance of anesthesia during surgery were achieved using SomnoSuite Low-Flow Anesthesia System (Kent Scientific Corporation). Following induction of anesthesia, animals were quickly secured to the stereotaxic apparatus (David KOPF instruments). Anesthesia depth was validated by toe-pinching and manual heart-rate monitoring. Isoflurane levels were adjusted (0.8–1.5%) to maintain a heart rate of ∼60 bpm. The skin was cleaned with Betadine (Dr. Fischer Medical); Lidocaine (Rafa Laboratories) was applied to minimize pain; and Viscotears gel (Bausch & Lomb) was applied to protect the eyes. An incision was made to expose the skull, which was immediately cleaned with hydrogen peroxide, and a small hole was drilled using a fine drill burr (model 78001RWD Life Science). Using a microsyringe (33GA; Hamilton syringe) connected to an UltraMicroPump (World Precision Instruments), a virus was subsequently injected at a flow rate of 50–100 nl/min, after which the microsyringe was left in the tissue for 5–10 min after the termination of the injection before being slowly retracted. A fiberoptic ferrule (400 μm, 0.37–0.48 NA, Doric Lenses) was slowly lowered into the brain. A custom-made metal head bar was glued to the skull, the incision was closed using Vetbond bioadhesive (3M), and the skull was covered in dental cement and let dry. An RFID (radio-frequency identification) chip (ID-20LA, ID Innovations) used for tracking during behavioral training was implanted subcutaneously. Mice were then disconnected from the anesthesia and were administered a subcutaneous saline injection for hydration and an IP injection of the analgesic Rimadyl (Norbrook) as they recovered under gentle heating. Coordinates for the claustrum were based on the Paxinos and Franklin mouse brain atlas (Paxinos & Franklin, 2013). retroAAV-CKII-iCre virus was prepared at the vector core facility of the Edmond and Lily Safra Center for Brain Sciences at the Hebrew University, as described previously (Atlan et al., 2017), and was injected to the coordinates ACC: ±0.25, 1.1, -1.75 mm relative to bregma, volume of injection : 200nl. AAV9-CAGGS-FLEX-ChR2-TdTomato was purchased from UPENN virus core facility, the coordinates of injection CLA: ±2.8, 1, -3.7; ±3.25, 0, -4.15 mm relative to bregma, volume of injection: 200nl. Surgeries for NPXL probe insertion are described below.

### Histology

Mice were anesthetized for terminal perfusion by a ketamine/xylazine mix and perfused with RT PBS, followed by cold 4% PFA. Following decapitation, heads were placed in 4% PFA overnight to preserve the location of the optic ferrule. Brains then were carefully extracted and placed in 4% PFA for another night prior to transitioning to PBS in preparation for sectioning and staining. The fixed tissue was sectioned using a Vibratome (Campden 7000smz-2) at 100 µm thickness.

### Image acquisition

Slides were scanned on a high-speed, fully-motorized, multi-channel epifluorescent light microscope (Olympus IX-81). Slices were imaged at 10X magnification (NA = 0.3). Green and red channels exposure times were selected for optimal clarity and were kept constant within each brain series. eGFP, excitation 490 ± 20 nm and emission 525 ± 36 nm and Alexa 647, excitation 625 nm and emission 670 nm.

### Histology registration and NPXL probe localization

Coronal sections were manually registered to the Allen Mouse Brain Common Coordinate Framework (CCF v3; 10 µm, 2017 release) (Wang et al., 2020) using MATLAB tools from the cortex-lab allenCCF repository (https://github.com/cortex-lab/allenCCF), with control-point warping to the template followed by reconstruction of Neuropixels probe trajectories and assignment of brain areas along the track. The workflow follows established pipelines for slice alignment and probe-track analysis (e.g., SHARP-Track and AP_histology) (Shamash et al., 2018) (https://github.com/petersaj/AP_histology).

#### ENGAGE behavioral task

Mice were initially trained in an automated behavioral cage system (see *Automated behavioral training* below) before undergoing habituation to head fixation and task performance on a low-friction, rodent-driven linear treadmill (Janelia 2017-049). The ENGAGE task (Fig. 1A) consisted of fixed-paced trials automatically initiated every 18 seconds. Behavioral sessions (total of 24 test sessions) consisted of 297 ± 36 trials. The task was controlled using a ‘Bpod’ state machine (Sanworks), with a custom MATLAB protocol.

Each trial began with a brief (100 ms) broadband noise stimulus (“Wait” signal), during which mice were required to withhold licking. The ‘Wait’ signal was followed, after a variable delay (0.75–2.5 s), by the ‘Go’ cue—five 6 kHz pure-tone pips (100 ms each). Licking during the delay period resulted in immediate trial termination and was classified as a *premature error*. After ‘Go’ cue onset, a 1.5 s response window was provided for a correct lick (*hit*), which was rewarded with a 4 µl drop of sucrose solution. Trials in which no lick occurred within this window were classified as *omissions* and were unrewarded.

Trial difficulty was modulated by two independent parameters: (1) four equally probable ‘Go’ cue intensities and (2) the presence or absence of a tone-cloud mask (50% of trials; 4 s of continuous chords assembled from logarithmically spaced pure tones in the range 1–10 kHz, excluding 6 kHz). The effect of cue intensity was quantified as Selectivity Index = (Hit rate @ highest intensity)−(Hit rate @ lowest intensity)

#### Optogenetic stimulation

We used a commercially available LED (Cat #LEDFLS_450, Doric LTD, Franquet, Quebec, Canada), emitting light at a wavelength of 450nm. We used a light intensity of 7mW per hemisphere. Optogenetic laser stimulation was delivered in one of three conditions: no stimulation (50% of trials), stimulation surrounding the ‘Wait’ signal (−0.5 s to +0.5 s relative to trial onset; 25% of trials), or stimulation surrounding—but not predictive of—the ‘Go’ cue (+0.5 s to +3 s after trial onset; 25% of trials).

#### Automated behavioral training

Mice were trained on the ENGAGE task using a custom, automated behavioral setup based on the *Bpod* system (Sanworks), modified to support group-housed training as described in (Peretz-Rivlin et al., 2024): The setup, built directly in the home cage, enables self-initiated, simultaneous, and individualized training of co-housed mice within their familiar environment.

Training cages consisted of a 4cm diameter tube corridor connected to the home cage, with a behavioral lick port (Sanworks) positioned at the far end. Food was available *ad libitum*, while water was provided exclusively via the behavioral system. An RFID reader (ID-20LA, ID Innovations) located above the corridor allowed automatic identification of individual mice. Auditory cues were delivered via a Bpod wave player (Sanworks) to earphones positioned adjacent to the lick port. Task control was implemented in MATLAB using the open-source Bpod state machine, integrated with RFID output to allow individualized training parameters based on each mouse’s performance.

Training began with a lick adaptation phase, in which mice learned to associate an auditory–visual cue with water delivery. Subsequent stages gradually increased task difficulty toward the full ENGAGE task as implemented under head-fixation for NPXL recordings and optogenetic stimulation (see *ENGAGE behavioral task* above). A trial was initiated when a mouse entered the port (RFID detection), immediately followed by the trial onset ‘wait’ signal. Premature or late licks were not rewarded, requiring the mouse to exit and re-enter the port in order to reinitiate a new trial.

**Stage 1**: Random delay 0–0.5 s (average 2.6 days to criterion; 50–70% correct).

**Stage 2**: Delay extended to 0.5–2 s (average 2.5 days).

**Stage 3**: Full delay range (0.5–3 s) and gradual removal of the visual cue in auditory trials: 30% Aud (Stage 3a), 50% Aud (3b), 70% Aud (3c) (average 4.4 days).

**Stage 4**: Introduction of tone-cloud masking stimulus (4 s of continuous chords, logarithmically spaced 1–10 kHz tones excluding the 6 kHz target frequency, 67.5 dB SPL).

**Stage 5**: Addition of three levels of target cue attenuation.

Across all stages, reward size (4 μl sucrose) and response window duration were constant. Total automated training time averaged 29.6 days.

#### In-vivo Electrophysiology Recording

On the first day of recording, mice were anesthetized using sevoflurane (Induction: 8%, Maintenance: 2-3%, SomnoSuite). A small craniotomy was then drilled (model 78001RWD Life Science) in a region of the skull approximately 0.5 to 1 mm anterior, and -0.3 to 0.3 mm lateral to Bregma. The craniotomy was covered with a silicone elastomer (WPI; Kwik-Cast cat#KWIK-CAST), and mice were allowed at least 30 minutes to recover from anesthesia in their homecage. Gentle heating was provided during the first few minutes of recovery. Following recovery, mice were head-fixed again. The silicone cap was removed, and an electrical ground (Ag/AgCl) was located on the surface of the skull and covered with saline. A Neuropixels probe (imec, phase 1.0) was lowered to the level of the skull at an angle of 20° relative to the anterior axis, and inserted through the craniotomy to a depth of 4.6 mm. Probes were covered with fluorescent dyes (DiI [Invitrogen cat#V22885], DiO [Invitrogen cat#V22886] or DiD [Invitrogen cat#V22887]) before penetration, to enable reconstruction of penetration sites. Following insertion, we waited 20 minutes to allow the tissue to stabilize around the probe. Then, we initiated data acquisition, and the ENGAGE behavioral session.

All recordings were acquired using Neuropixels phase 1.0 probes (Imec) (Jun et al., 2017) connected to an Imec base-station and an National Instruments PXIe-6341 auxiliary board. Signals were sampled at 30 kHz. The action potential (AP) band was filtered between 0.3–10 kHz, and the local field potential (LFP) band between 0.1–1000 Hz (sampled in 2.5 kHz). Data acquisition was performed with **SpikeGLX** software (Release_v20202203-phase30; https://billkarsh.github.io/SpikeGLX/). All the task parameters timings were recorded simultaneously with the auxiliary NI board.

Preprocessing was carried out using the ‘CatGT’ tool (https://billkarsh.github.io/SpikeGLX/#catgt). Steps included *t_shift* correction, band-pass filtering of the AP stream (300–9000 Hz) and LFP stream (0.1–1000 Hz), application of a global common-average reference (CAR) filter across AP channels, and artifact removal using *gfix*. Behavioral event times were aligned to the Neuropixels data stream using ‘<u>TPrime’</u> (https://billkarsh.github.io/SpikeGLX/#tprime).

#### Spike sorting

Preprocessed electrophysiological data were spike-sorted using Kilosort 3 (Pachitariu et al., 2024). Sorted units were then manually curated in **Phy2** (Template GUI, https://github.com/cortex-lab/phy ), including the merging of clusters corresponding to the same unit and removal of artifacts or noise clusters. After manual curation, we applied the ‘remove_duplicates’ function from the Allen Institute’s open-source spike-sorting pipeline (Allen Institute, *ecephys_spike_sorting*; https://github.com/AllenInstitute/ecephys_spike_sorting) to eliminate duplicate spike detections across neighboring channels. To ensure reliability, we then applied a spike-amplitude threshold tailored to each recording based on its signal-to-noise ratio (SNR), retaining only high-SNR units with stable waveforms. Finally, only units with an average firing rate greater than 0.1 Hz across the session were included in subsequent analysis.

#### Single-unit responses and clustering

For each recorded unit, spike times were aligned to task events, defined according to trial outcome: *correct trials* (Wait signal, Go cue, Rewarded lick), *premature errors* (Wait signal, Unrewarded lick), and *omissions* (Wait signal, Go cue). Peri-event raster plots and peri-stimulus time histograms (PSTHs) were generated for each unit. Spikes were counted in 20 ms bins relative to the event, divided by the number of event repetitions, and smoothed with a three-bin running average.

To normalize responses across units, each PSTH was baseline-corrected: the mean firing rate during a 2 s pre-trial window (averaged across trials) was subtracted from the PSTH and the result was then divided by this baseline, allowing comparison across units with different spontaneous firing rates. All traces and heatmaps shown in the figures are presented in baseline-normalized (non–z-scored) units.

Preprocessing for clustering and ordering involved additional concatenation of all baseline-normalized PSTHs for a given event and unit were into a matrix of dimensions *n* units × *n* time bins. This matrix was z-scored along rows (time bins within each unit. The z-scored matrix was input to the **Rastermap** algorithm (Stringer et al., 2025) https://github.com/MouseLand/rastermap ) with the following parameters: *n_clusters* = 18, *n_PCs* = 50, *locality* = 0.5, *time_lag_window* = 15, *grid_upsample* = 10, *bin_size* = 1. Concatenated PSTHs were plotted according to the Rastermap embedding order (Figures 2–3). Units were assigned to clusters based on rounding their embedding values, yielding 19 clusters in total. For each cluster, concatenated PSTHs were averaged across units, and hierarchical clustering (median linkage, cityblock distance) was applied to group clusters into higher-order categories.

#### Population decoding

To decode behavioral outcomes from neural activity, population firing-rate vectors were constructed in 100 ms bins within defined windows around the Go cue. For each time bin, balanced datasets (equal numbers of trials per outcome) were generated by bootstrap resampling, repeated 1000 times. On each resampled dataset, a logistic regression classifier was trained to discriminate between two trial outcomes based on the population activity vectors. Decoding accuracy was averaged across repetitions to yield per-session performance estimates.

#### CEBRA embedding

To characterize neural population dynamics and their relationship to task epochs, we used CEBRA (Schneider et al., 2023) , a supervised dimensionality reduction method that embeds neural activity into a low-dimensional space while preserving task-relevant structure.

For each unit included in the analysis, firing rates were computed in 50 ms bins across the session. Each time bin was labeled according to the corresponding task epoch: inter-trial interval (ITI); *Wait* period (from Wait signal to Go cue in correct trials, from Wait to unrewarded lick in premature errors, and from Wait to Go cue in omissions); *Premotor* (Go cue to rewarded lick in correct trials); *Reward feedback* (3 s following reward delivery); *Premature feedback* (3 s following an unrewarded lick); and *Omission feedback* (3 s following an omitted Go cue).

A CEBRA model https://github.com/AdaptiveMotorControlLab/CEBRA , was trained in supervised mode using these discrete labels, separately for ChR2 sessions and DIO-TdTomato sessions (multisession training). Model parameters were: *model_architecture* = “offset10-model”, *batch_size* = 1024, *temperature_mode* = “auto”, *temperature* = 1, *output_dimension* = 3, *max_iterations* = 60,000, *distance* = “cosine”, *conditional* = “time_delta”, and *time_offset* = 10.

Decoding analyses were performed per session using subsets of the embedding restricted to the labels of interest. For each subset, balanced datasets (equal numbers of points per label) were generated by random sampling, repeated 1000 times. On each sampled dataset, a *k*-nearest neighbors (KNN) decoder (*k* = 10) was trained using the CEBRA package, and decoding accuracy was averaged across repetitions to yield a per-session value.

#### Local field potential analysis

For each recording session, 10 adjacent channels localized to the ACC (based on reconstructed probe trajectories) were selected. The median across these channels was calculated to obtain a robust, noise-reduced LFP signal. This signal was bandpass-filtered into two frequency ranges of interest: delta (0.5–4 Hz) and theta (6–12 Hz). Band-specific power was then computed by applying the Hilbert transform to the filtered signal. To allow comparisons across sessions, power values were normalized by z-scoring over the entire session. Finally, for each frequency band, power time courses were aligned to task events and averaged across repetitions of each event type.

#### Code and figure preparation

All analyses were performed using custom-written code in Python (packages including NumPy, SciPy, scikit-learn, pandas, xarray, and Matplotlib). Figures were generated in Python and subsequently formatted and annotated in Adobe Illustrator for presentation.

## Code and Data availability

Code and data will be made available upon reasonable request from the corresponding author

---

**Supplementary Figure 1:**
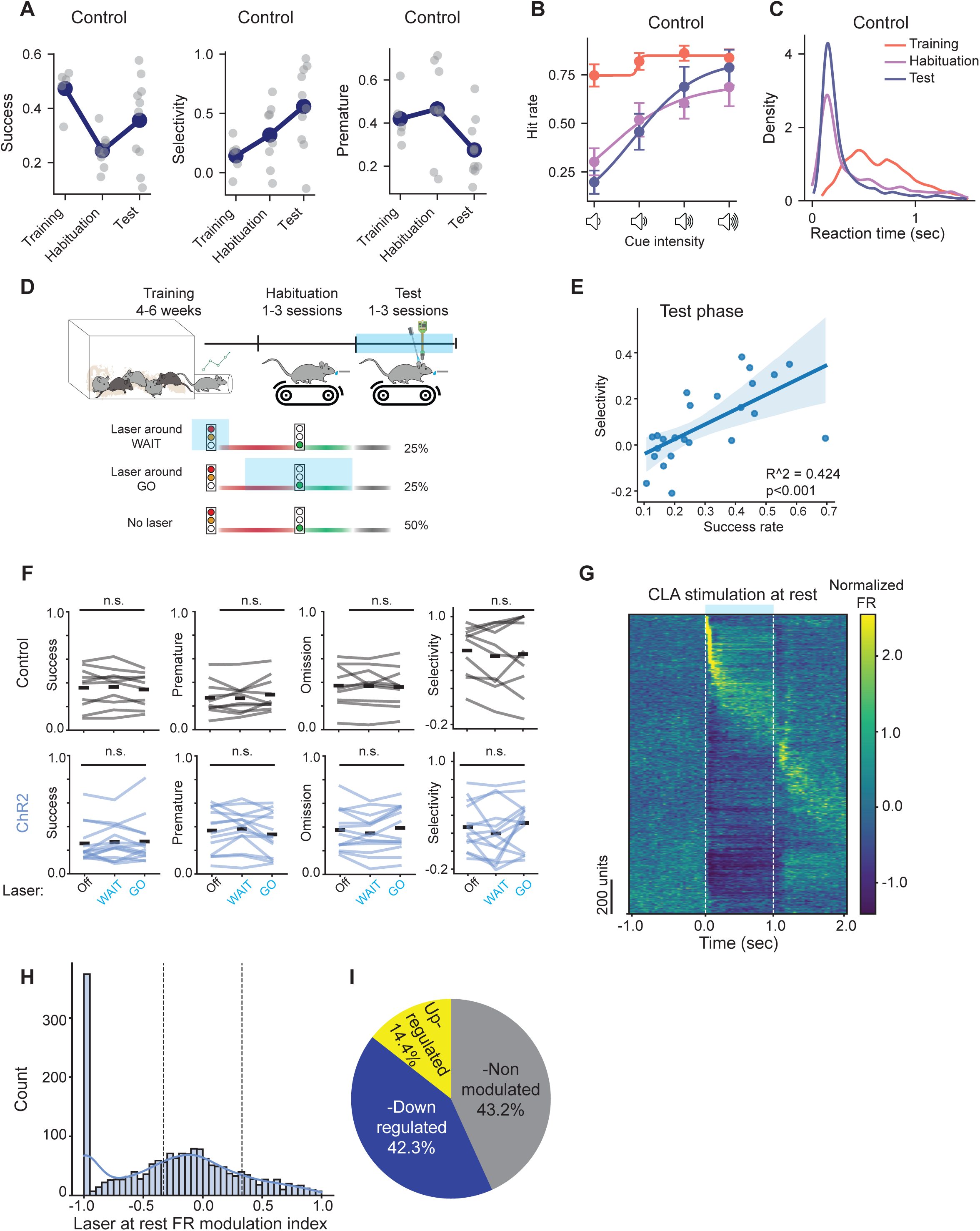
Behavior and Validation of Optogenetic Manipulation. (**A**) Longitudinal performance for control mice across task phases. Home cage points represent individual animal averages over the last 1000 trials of automated training (n = 5 mice); Head-fixed points represent individual sessions (Habituation: n = 8 sessions from 5 mice; Test: n = 10 sessions from 5 mice). Left: success rate, middle: Selectivity Index (defined as the difference in hit rate between highest and lowest intensity cues), right: premature error rate. (**B**) Psychometric curve of hit rate vs. GO cue sound intensity in the three phases of the experimental timeline. Points indicate averages, error bars correspond to SEM (training n=5 mice, habituation n = 8 sessions, test n = 10 sessions). (**C**) Distribution of reaction times in Hit trials of control mice in different task phases – Training, Habituation and Test (**D**) Experimental timeline. Top: Mice were first trained in an automated homecage system for 4–6 days to acquire task contingencies, and then transferred to a head-fixed setup for recording. After 1–3 habituation sessions, mice underwent 1–3 test sessions with concurrent optogenetic stimulation and Neuropixels recordings. Bottom: Laser stimulation was delivered in three trial types: in 50% of trials, no stimulation was applied; in 25% of trials, stimulation was aligned to the ’Wait’ signal (-0.5 to +0.5 s relative to onset); and in the remaining 25%, stimulation occurred 0.5–2.5 s after the ’Wait’ signal to overlap with the ’Go’ cue without providing a predictive timing cue. (**E**) Correla-tion between selectivity and success rates, points indicate individual sessions from the test phase (including controls and ChR2, n = 24 sessions; R2=0.424, p<0.001). (**F**) No acute behavioral effects of laser stimulation were observed within sessions. Success rate, premature error rate, omission rate, and selectivity index (high–low intensity hit rate difference) did not differ significantly across stimulation conditions. Mixed-effects linear model: behavior ∼ laser + (1|session). Top: Control (n = 10 sessions); Bottom: DIO-ChR2 (n = 14 sessions). (**G**) Heatmap of normalized firing rates for frontal cortex units in response to 1 s claustrum laser stimulation delivered during rest (n= 1917 units) (H) Distribution of significantly up- and down-regulated frontal units in response to claustrum stimulation. (**I**) Distribution of laser modulation index values.

**Supplementary Figure 2.**
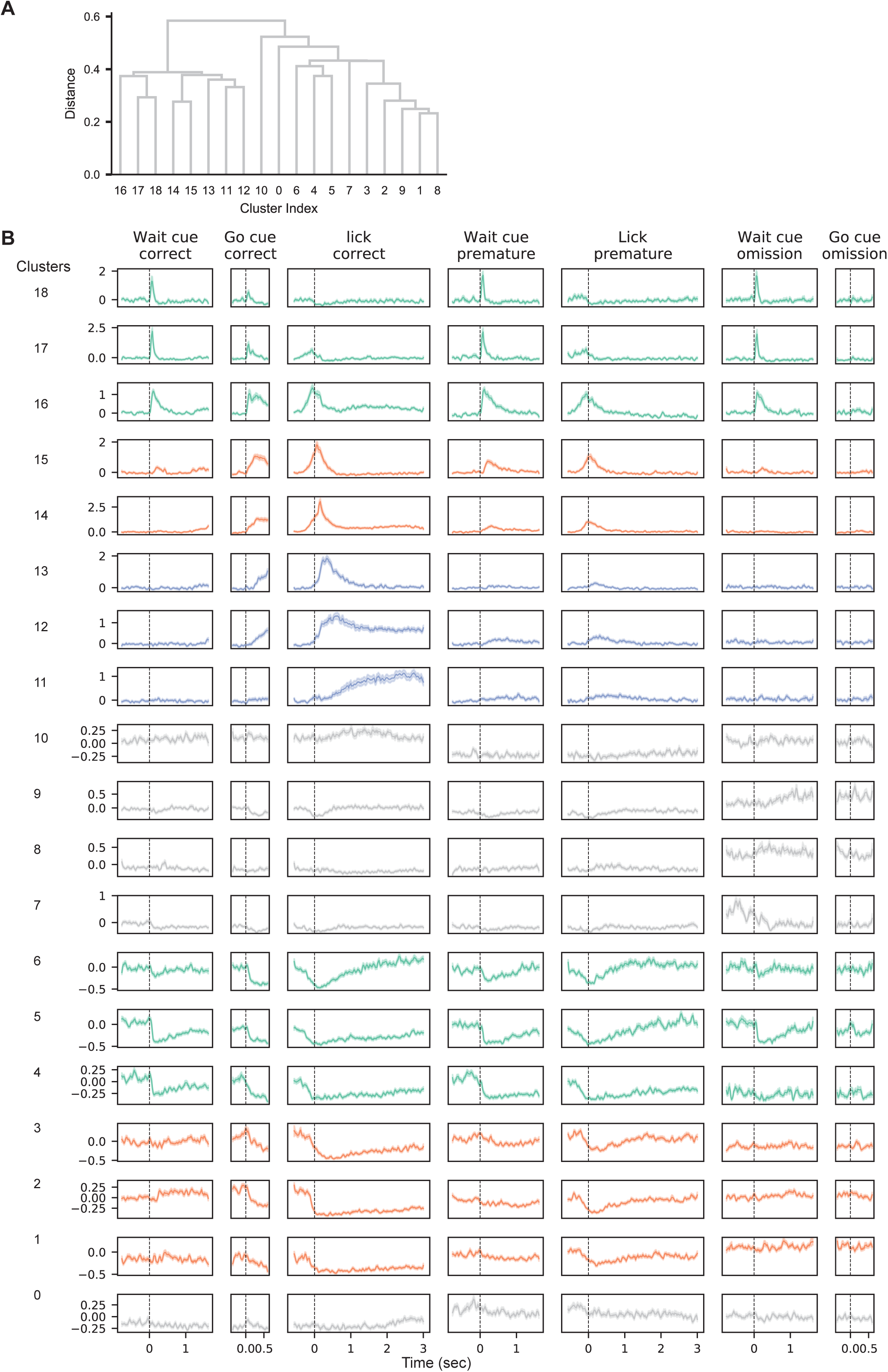
Functional classification of frontal cortex neurons and extended response profiles across task events. (**A**) Hierarchical clustering of the 19 clusters generated by the Rastermap algorithm. Clusters were grouped based on similarity of average temporal response profiles across task events using Median on cityblock distances of mean PSTHs. (**B**) Average PSTH for each of the 19 clusters generated by Rastermap, aligned to all task events: correct trials (WAIT signal, GO-cue, rewarded lick), premature trials (WAIT signal, unrewarded lick), and omission trials (WAIT signal, GO-cue). Lines represent means; shaded areas indicate SEM. Clusters are color-coded by functional classification as shown in Figure 2B.

**Supplementary Figure 3.**
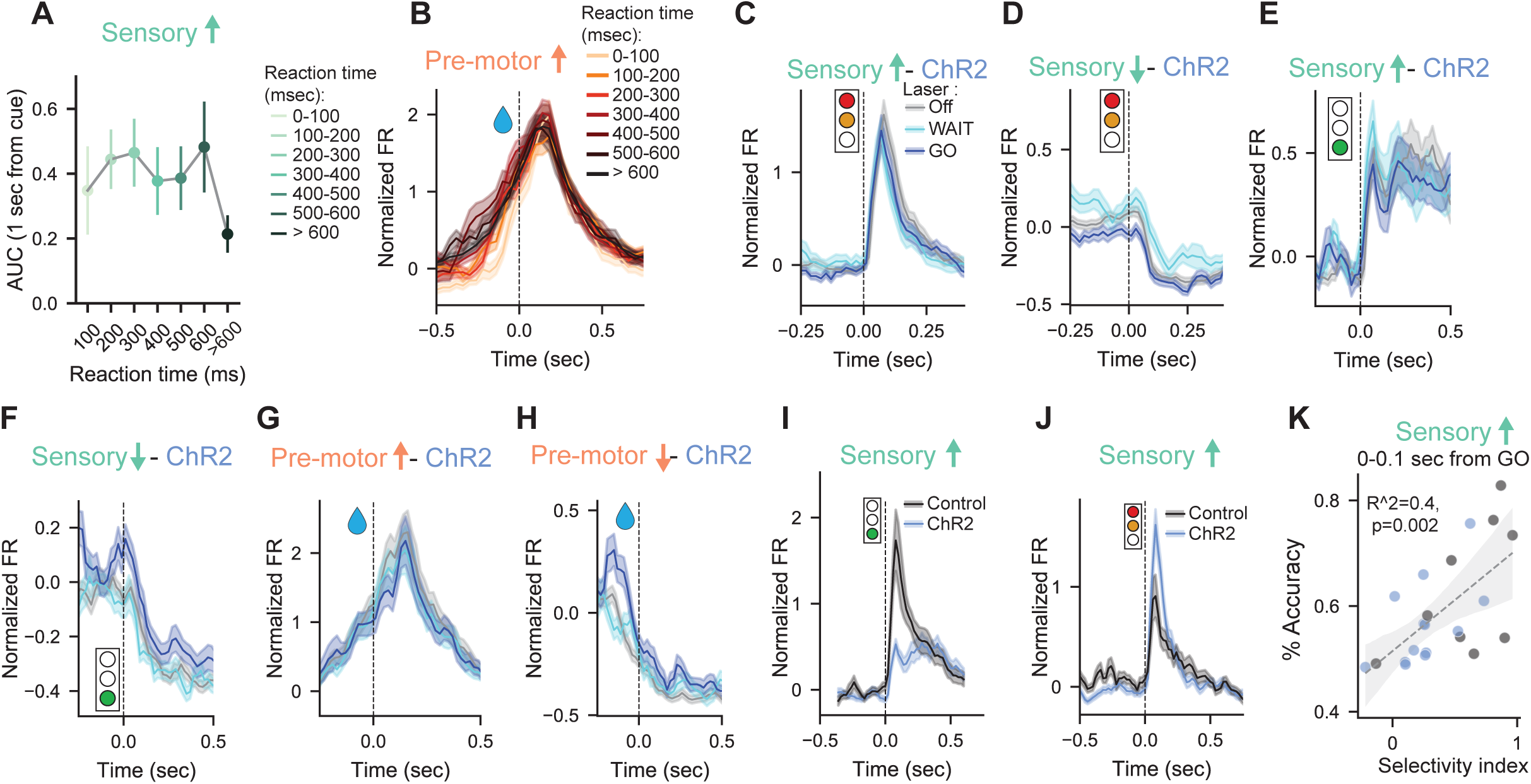
(**A**) Area under the curve (AUC) of normalized firing rate in the first 1000 ms following GO-cue onset in hit trials, plotted against reaction time bins. Points represent means; error bars indicate SEM (n = 66 units). (**B**) PSTH of premotor-upregulated units from control mice (n = 115) to lick action in hit trials, separated by reaction time quantiles. (**C-H**) Responses of recorded units from ChR2 mice were not significantly altered by laser condition (no laser, laser during WAIT, laser during GO). Lines show means; shaded areas, SEM. (**C**) Sensory-upregulated units aligned to the WAIT signal (n = 127). (**D**) Sensory-downregulated units aligned to the WAIT signal (n = 229). (**E**) Sensory-upregulated units aligned to the GO-cue in hit trials (n = 127). (**F**) Sensory-downregulated units aligned to the GO-cue in hit trials (n = 229). (**G**) Premotor-upregulated units aligned to rewarded licks (n = 115). (**H**) Premotor-downregulated units aligned to rewarded licks (n = 294). (**I**) Average activity triggered by the GO-cue in hit trials comparing recordings from Controls to ChR2 mice, for sensory-upregulated units (66 vs. 127 units) (**J**) Average activity triggered by the WAIT-cue in hit trials comparing recordings from Controls to ChR2 mice, for sensory-upregulated units (66 vs. 127 units). Lines show means; shaded areas, SEM. (K) Decoding performance for predicting hit vs. omission trials following ‘Go’ cues from early (0–100 ms) sensory-upregulated responses versus behavioral selectivity (R² = 0.40, p = 0.002).

